# Mutations in conserved residues of the myosin chaperone UNC-45 result in both reduced stability and chaperoning activity

**DOI:** 10.1101/2021.05.07.443121

**Authors:** Taylor Moncrief, Courtney J. Matheny, Ivana Gaziova, John Miller, Hiroshi Qadota, Guy M. Benian, Andres F. Oberhauser

## Abstract

Proper muscle development and function depends on myosin being properly folded and integrated into the thick filament structure. For this to occur the myosin chaperone UNC-45, or UNC-45B, must be present and able to chaperone myosin. Here we use a combination of *in vivo C. elegans* experiments and *in vitro* biophysical experiments to analyze the effects of six missense mutations in conserved regions of UNC-45/UNC-45B. We found that the phenotype of paralysis and disorganized thick filaments in 5/6 of the mutant nematode strains can likely be attributed to both reduced steady state UNC-45 protein levels and reduced chaperone activity. Interestingly, the biophysical assays performed on purified proteins show that all of the mutations result in reduced myosin chaperone activity but not overall protein stability. This suggests that these mutations only cause protein instability in the *in vivo* setting and that these conserved regions may be involved in UNC-45 protein stability/ regulation via post translational modifications, protein-protein interactions, or some other unknown mechanism.

## INTRODUCTION

Multiple studies have demonstrated that myosin heads do not fold spontaneously and likely require chaperones (1–3).The first myosin head chaperone discovered was UNC-45, from studies in *C. elegans*. UNC-45 homologs are found in all metazoa (4). Drosophila and *C. elegans* have one UNC-45 gene expressed in all cells, whereas vertebrates have two UNC-45 genes, one expressed in striated muscle (UNC-45B) and one expressed in all cells (UNC-45A). The first clue to the existence of UNC-45 came from classical genetics experiments in *C. elegans*. The gene was identified by a temperature sensitive mutant, *unc-45(e286)*, that when grown at the restrictive temperature of 25° shows striated muscle with disorganized sarcomeres and reduced numbers of thick filaments (5,6). Identification of the gene at the molecular level showed that it encodes a protein of 961 amino acids and comprised of 3 regions: a 100 residue long TPR (tetratricopeptide repeat) region, a 400 residue central region, and a 430 residue UCS (UNC-45, CRO1, She4p) domain (6,7). The UCS domain was defined by sequence similarity to two fungal proteins, CRO1 and She4p, that also functionally interact with myosin. When grown at the restrictive temperature, *unc-45(e286)* animals show decreased accumulation of MHC B (myosin heavy chain B), the major myosin heavy chain of thick filaments in the body wall muscle of C. elegans (6). These thick filaments consist of MHC A in a small middle portion, and MHC B in the major outer portions (8). Interestingly, antibodies to UNC-45 co-localize with MHC B but not with MHC A in sarcomeric A-bands in already assembled sarcomeres of adult muscle (9,10).

The regions or domains of UNC-45 are functionally distinct. The TPR region binds to heat shock protein 90 (Hsp90), and the central region and UCS domain bind to myosin heads (11). UNC-45 can inhibit the thermal aggregation of myosin heads in vitro, suggesting that indeed UNC-45 chaperones the myosin head (11). *unc-45* mutants can be rescued by full length UNC-45, UNC-45 in which the TPR has been deleted, and even the UCS domain itself (12). The crystal structure of *C. elegans* UNC-45 (10) shows that the overall shape is of an “L” with the UCS domain forming one leg, and the central and TPR forming the other leg. In addition, all of the central region and UCS domain consists of 17 armadillo (ARM) repeats, each consisting of 2-3 α-helices. Most significantly, UNC-45 forms linear multimers in which the length of the repeating unit (17 nm) is similar to the repeating unit in the staggered display of pairs of myosin heads along the surface of thick filaments (closest distance between adjacent double heads is 14.3 nm). Thus, these UNC-45 multimers may help stabilize this arrangement during thick filament and sarcomere assembly. Also, this architecture suggests the possibility that in the assembled thick filament in which UNC-45 seems to be associated, may aid refolding of the myosin heads that are damaged by heat or oxidation as the muscle is used and ages.

Based upon actin filament gliding assays of myosin heads with and without the presence of UNC-45 and/or HSP-90 (13,14), the following model can be envisioned: Under normal conditions the UCS domain of UNC 45 is bound to the myosin head and the TPR domain is bound to HSP 90, and yet the myosin head is active in pulling actin thin filaments. However, under stress conditions, HSP 90 detaches from the TPR domain, causing a conformational change in UNC-45 that allows the central domain to bind to the myosin neck resulting in inhibition of the myosin power stroke while the UCS domain re-folds the myosin head. Then, when the refolding is complete, some sort of signal is received for HSP-90 to re-bind the TPR domain of UNC-45, causing the central domain to release the myosin neck, thus allowing the myosin power stroke to resume.

Two major phenotypic classes of mutants affect *C. elegans* sarcomere assembly and muscle activity (15): (1) A subclass of the “uncoordinated” (Unc) class which are viable adults that are either paralyzed or move more slowly or in a strange way compared with wild type; although most of the 129 Unc genes affect the nervous system, 40 are specific for muscle. (2) The “paralyzed arrested at two-fold” (Pat) embryonic lethal class of 20 genes. These genes encode proteins essential for embryonic sarcomere assembly, especially components of the integrin adhesion complexes. For some genes, the null state is Pat, and the loss of function or hypomorphic state is Unc, and *unc-45* is an example of such a gene. For *unc-45*, the Pat embryonic lethal mutants have stop codons upstream of the UCS domain, whereas the Unc mutants, all of which are temperature sensitive (*ts*), are amino acid substitutions in the central and UCS domains (6,7,16,17).

In this report, we first expanded the number of known *C. elegans unc-45* Unc alleles by two, assessed the effect of all 6 known missense mutations on sarcomere organization and measured the levels of UNC-45 mutant proteins as compared to wild type UNC-45 protein. We then created the same mutations in the comparable residues of human UNC-45B, expressed and purified the proteins in *E. coli*, and characterized their thermal stability, and myosin chaperone activities. Through a combination of *in vitro* and *in vivo* data, we can conclude that the unc (paralysis and disorganized thick filaments) phenotype is caused by both a decline in chaperoning activity and a decline in overall steady state protein levels.

## RESULTS

### Effects of *unc-45* temperature-sensitive mutants on sarcomere organization, MHC B and UNC-45 protein levels

There are four well known temperature-sensitive (ts) mutations in *C. elegans* UNC-45: one found in the central domain (G427E) and three found in the UCS domain (L559S, E781K, L822F). To obtain additional ts mutants in the UNC-45 UCS domain, we used WormBase to look for missense mutations in conserved residues of the UCS domain amongst the Million Mutation Project (18) mutant strain collection. Six strains conforming to these criteria were obtained from the Caenorhabditis Genetics Center and were grown at the permissive temperature of 15°, and the restrictive temperature of 25°, and immunostained with antibodies to myosin heavy chain A in order to assess the organization of assembled thick filaments (A-bands) in adult muscle. From this analysis, we identified two new ts mutants, that showed normal or nearly normal thick filament organization at 15°, but disorganized thick filaments at 25°, and these are G703R and F724S. These mutant strains retained the same phenotype after outcrossing 5 times to wild type to remove most of the background mutations (data not shown). **Figure 1A** shows thick filament organization for the now new total of 6 ts missense mutations in the UCS domain of UNC-45.

**Figure 1.**
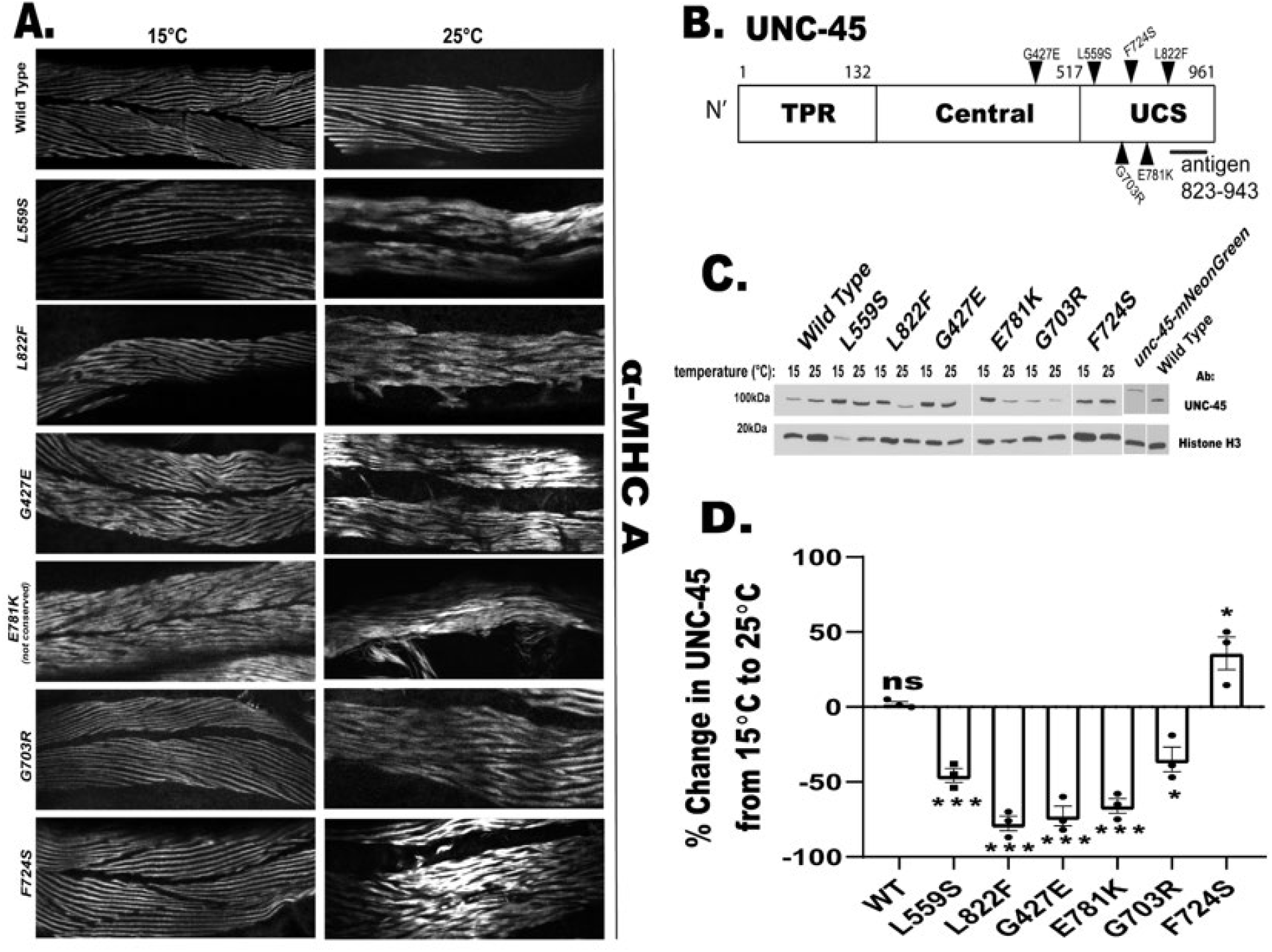
Effects of *unc-45* temperature-sensitive (ts) mutants on sarcomere organization and UNC-45 protein levels. **A)** Immunostaining of A-bands with antibodies to MHC A for wild type and the 6 *unc-45* mutant strains raised at 15° and 25°. For simplicity only the amino acid changes in the mutant UNC-45 proteins are used to distinguish the mutants (see Table 1 for mutant allele names). Except for wild type, all mutants showed more severe disorganization of sarcomeric A-bands at 25°. Three alleles, with mutations in F724S, G703R, and L559S, show normal organization of A-bands at 15°. **B)** Schematic of UNC-45 domain organization, location of 6 ts mutants examined, and the location of the antigen used to raise rabbit polyclonal antibodies to UNC-45. **C)** Representative western blots examining the levels of UNC-45 in wild type and the 6 ts *unc-45* mutants at permissive (15°) and restrictive (25°) temperatures. The second from the last lane contains an extract from *unc-45(syb789)*, which is a CRISPR engineered strain that expresses UNC-45 with an mNeonGreen tag fused to the C-terminus of UNC-45. Note the shift in size of this band (134 kDa) compared to the size (107 kDa) of UNC-45 bands from wild type and the mutants demonstrating the specificity of the antibody. **D)** Graphical representation of the % change in UNC-45 levels of *unc-45* mutant alleles when shifted from 15° to 25°.

**Table 1.** Summary of the impact of ts mutations on the biophysical properties of human UNC-45 chaperone and on *C. elegans* protein levels.

| Human UNC-45 ts-mutant name | *C. elegans* ts-mutant (allele) | Location of the mutation (ARM repeat) | Protein stability CD /SYPRO (Tm) | ATPase protection (% relative to WT) | %change in UNC-45 from 15 to 25°C |
| --- | --- | --- | --- | --- | --- |
| UNC45B -WT | WT |  | 39°C /46°C | 100 | 0 |
| UNC45B-G408E | G427E (b131) | highly conserved aa in sequence between 6H3 and 7H1 | 40°C /47°C | 80 | -70 |
| UNC45B-L540S | L559S (su2002) | Highly conserved aa in in loop between 9H3 and 10H1 | 41°C /47°C | 75 | -50 |
| UNC45B -G690R | G703R (gk576605)* | highly conserved aa in 12H3 | 38°C /45°C | 67 | -20 |
| UNC45B -F711S | F724S (gk615477)* | highly conserved aa in loop between 12H3 and 13H2 | 36°C/ 43°C | 73 | +30 |
| UNC45B -E767K | E781K (m94) | highly conserved aa in 14H2 | 36°C/ 43°C | 76 | -70 |
| UNC45B -L808F | L822F (e286) | conserved aa in 15H2 (F in Dm) | 40°C/ 47°C | 75 | -75 |

\*: new mutations obtained from the million mutation project

One interpretation of the reduced thick filament assembly in these mutants is that at the restrictive temperature their UCS domains display reduced myosin chaperone activity. However, an additional contribution to the phenotype might be that these missense mutations result in reduced levels of mutant UNC-45 proteins due to instability and degradation. To explore this possibility we wished to determine the steady state levels of these mutant UNC-45 proteins in comparison with the steady state level of wild type UNC-45 protein. Although antibodies to UNC-45 have been described by several labs (9,10) we wanted to generate our own antibodies for this and future studies. As shown in **Figure 1B**, we used a 120 residue segment of the UCS domain as antigen to generate rabbit polyclonal antibodies. After affinity purification, these antibodies reacted to the expected sized protein of ~100 kDa from wild type and a larger fusion protein from a CRISPR strain in which mNeonGreen was fused to the C-terminus of UNC-45. We then compared the levels of UNC-45 protein from wild type and the 6 ts mutant alleles at 15° and 25° (**Figure 1C**). As indicated in **Figure 1D**, all of the mutants except for F724S, showed a significant decrease in the level of UNC-45 protein when the temperature was changed from 15° to 25° degrees. This result indicates that some of the defect in thick filament assembly is due to reduced levels of UNC-45 protein, or in other words, not all of the phenotype can be attributed to reduced UCS activity.

We also measured the steady state protein level of the main client of UNC-45, body wall muscle myosin heavy chain B (MHC B), because it was reported that unc-45(e286) [L822F] has decreased MHC B, when this mutant is grown at the restrictive temperature (6,12,19). In **Figure 2**, we show that 4/6 unc-45 mutants have significantly reduced levels of MHC B at the restrictive temperature (25°C) compared to the permissive temperature (15°C). There was a decline in MHC B in the canonical *e286* [L822F] mutant, however it was not significant due to the levels of MHC B being already diminished at the permissive temperature. Consistent with this diminished level of MHC B at the permissive temperature, even at the permissive temperature, *e286* [L822F] shows some disorganization of A-bands (**Figure 1A**). The newly identified mutant F724S, which had increased UNC-45 at the restrictive temperature, also had increased MHC B at the restrictive temperature. We can see substantial myosin aggregation and disorganization in the A-band staining of this mutant, providing further evidence that too much UNC-45 also results in the Unc phenotype. Interestingly, we also observed about a 25% loss of MHC B in the wild type strain grown at 25°C (**Figure 2B**), despite there being no significant decline in the level of UNC-45 (**Figure 1D**) or in A-band organization (**Figure 1A**) at this temperature.

**Figure 2.**
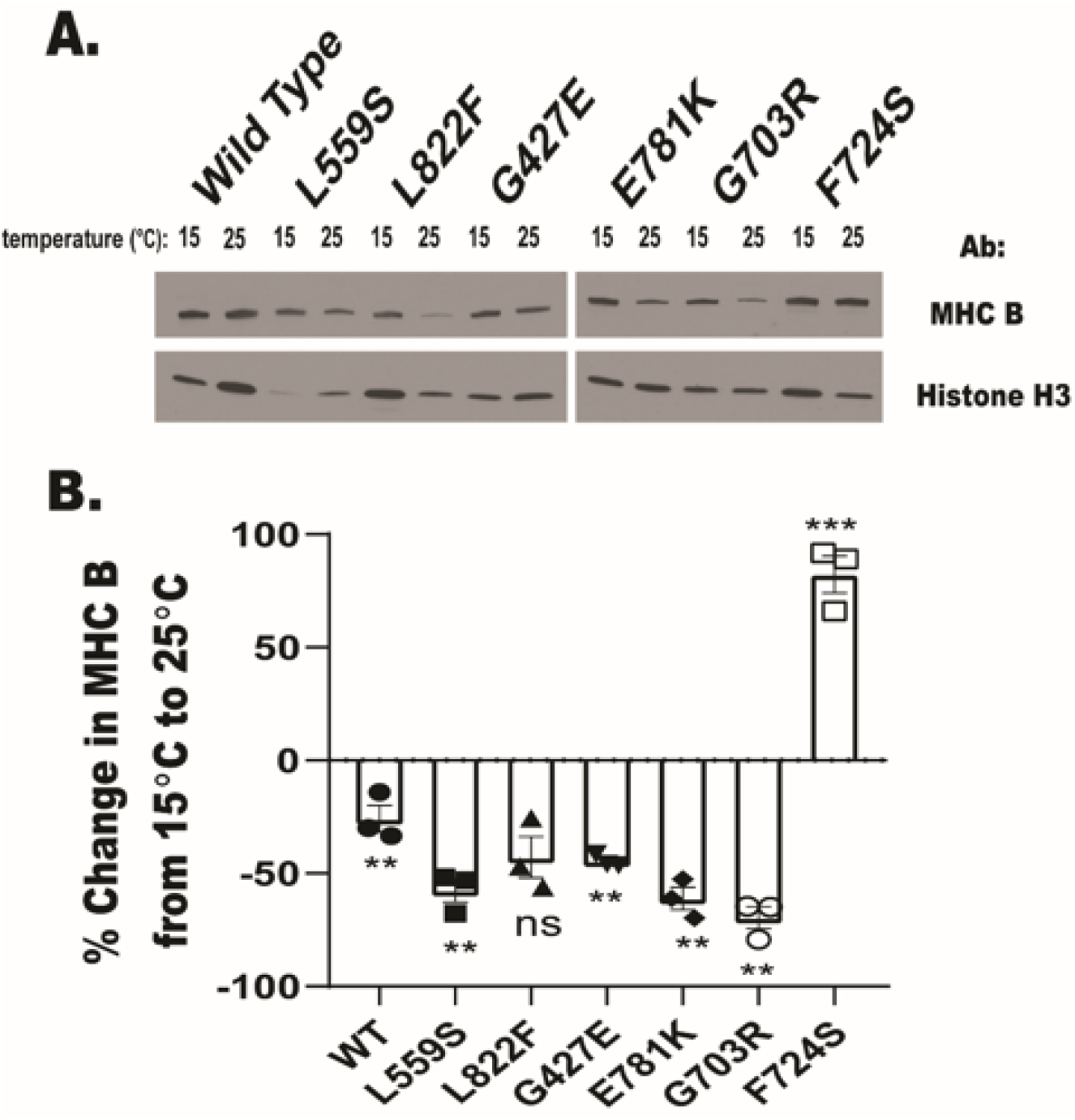
Effects of *unc-45* temperature-sensitive (ts) mutants on myosin MHC B protein levels. **A)** Representative western blots examining the levels of myosin MHC B in wild type and the 6 ts *unc-45* mutants at permissive (15°) and restrictive (25°) temperatures. **B)** Graphical representation of the % change in MHC B levels of *unc-45* mutant alleles when shifted from 15° to 25°.

### Temperature-sensitive mutations impact conserved residues in UNC-45B

In order to analyze the impact of these mutations on the structural stability and function of the UNC-45 chaperone, we introduced the equivalent mutations in the human UNC-45B sequence, because we found that wild-type and a wide variety of mutant proteins express well and as soluble proteins in *E. coli* (20), but this is not true for many mutant *C. elegans* UNC-45 proteins (unpub. data). The UNC-45B amino acid sequence is highly conserved down to invertebrates (10,17), which allowed us to explore the structural impacts of the ts-mutants on UNC-45B. A sequence alignment of UNC-45 proteins from *Homo sapiens* (Hs; Q8IWX7), *Drosophila melanogaster* (Dm; Q9VHW4) and *C. elegans* (Ce; Q09981) show that all six mutations are highly conserved (red residues are strictly conserved in **Figure 3**). Interestingly, the L882F mutation is found in the wild-type sequence of Drosophila UNC-45 protein, indicating that this mutation is structurally tolerated.

**Figure 3.**
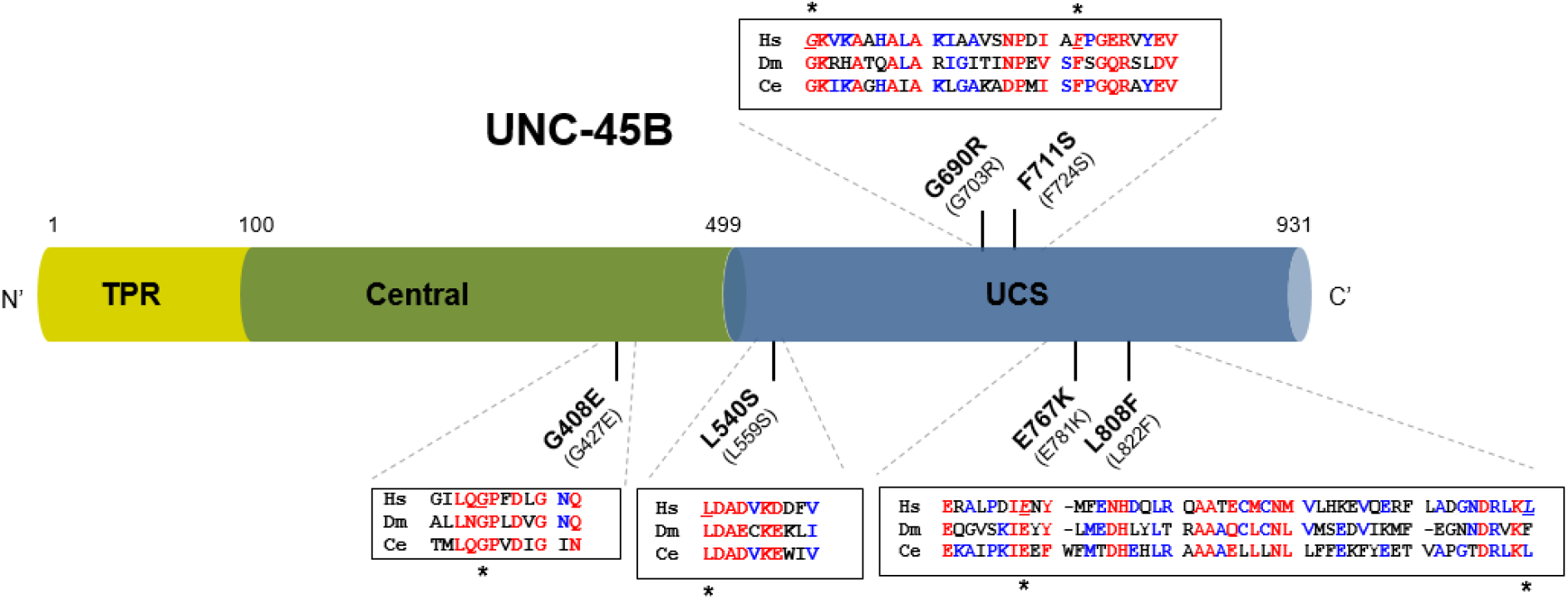
Location of the temperature-sensitive mutations in the human UNC-45B sequence. Human UNC-45B is made of three distinct domains: a 100-amino-acid N-terminal TPR domain (in yellow), a 400-amino-acid Central domain (in green), and a 431-amino-acid UCS domain (in blue). The equivalent *ts* mutations in the *C. elegans* UNC-45 sequence are shown in parenthesis. **Insets)** Sequence alignment of UNC-45 proteins from *Homo sapiens* (Hs; Q8IWX7), *Drosophila melanogaster* (Dm; Q9VHW4) and *C. elegans* (Ce; Q09981). Red residues are strictly conserved in all three sequences; blue are lower consensus residues. ClustalOmega (33) and Multalin (34) were used for the alignment. Mutated residues are marked by an asterisk (*) and underlined.

Although no human UNC-45 structure is available, homology models have been created via in silico molecular modeling (20,21). **Figure 4** shows a model of the human UNC-45B structure using the *C. elegans* crystal structure as a reference (PDB: 6qdl) (10,17). The insets in Figure 3 show the location of the six ts-mutations in the human structure. For example, G408E (G427E in *C. elegans*) lies in a loop between armadillo (ARM) repeats 6 and 7 (α-helices 6H3 and 7H1) and the introduced mutation does not seem to drastically interfere with the surrounding structures, according to the program Chimera calculations of van der Waals overlaps (22). All of the other mutations are predicted to be partially solvent exposed and hence do not sterically interfere with surrounding ARM repeats.

**Figure 4.**
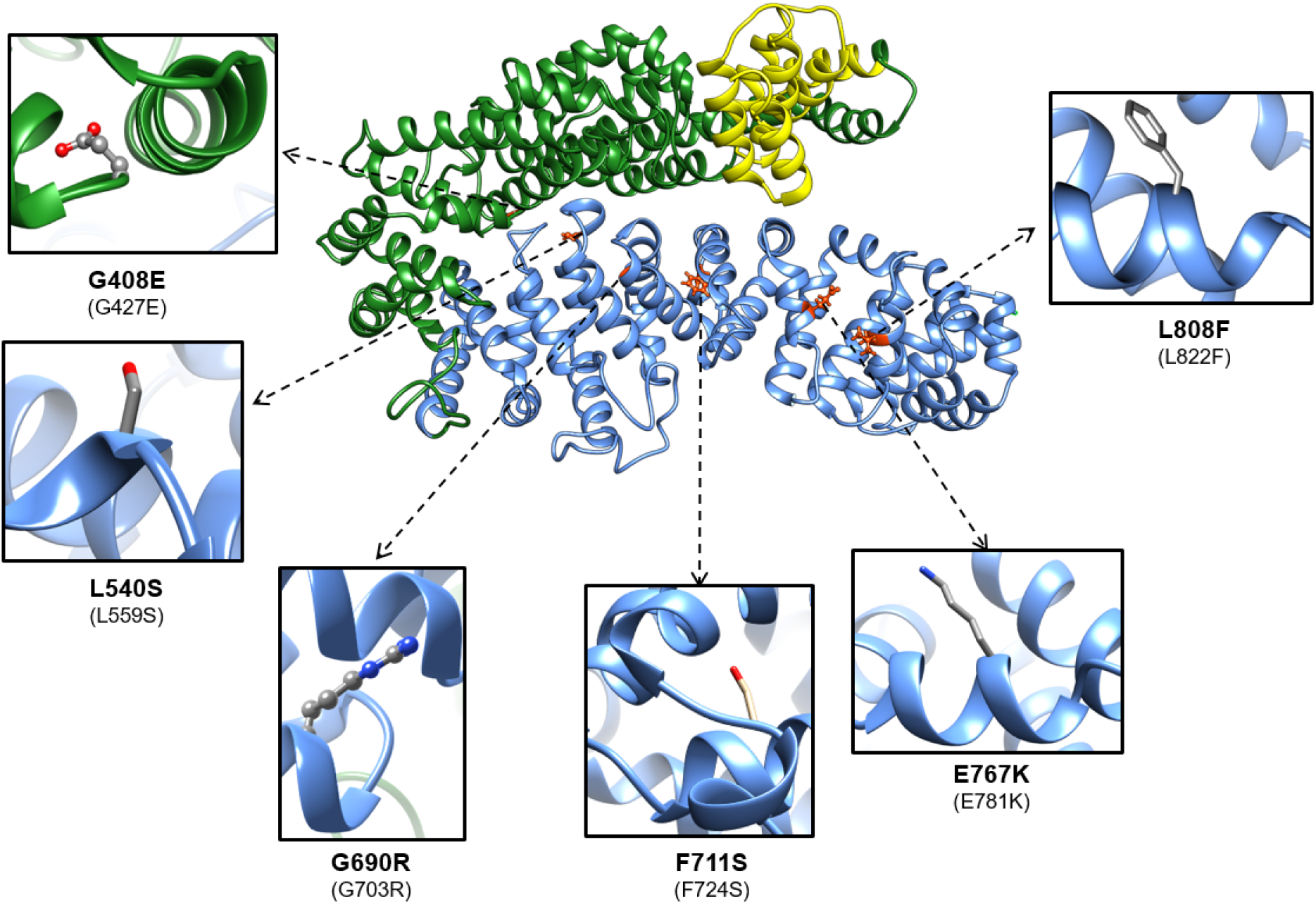
Location of the temperature-sensitive mutations in the human UNC-45B model structure. The homology model of the human UNC-45 protein structure was done using Phyre2 (30) and SWISS-MODEL (31) domain using the *C. elegans* 3D structure (PDB ID: 4i2z) as a reference crystal structure. The three domains are colored yellow (TPR), green (central) and blue (UCS). The corresponding temperature-sensitive residues were mutated in Chimera (22). The equivalent *ts* mutations in the *C. elegans* UNC-45 sequence are shown in parenthesis.

### Effects of temperature-sensitive mutations on thermal stability of human UNC-45B

To assess the impact of the mutations on the overall stability of the UCS domain we measured the changes in secondary structure as a function of temperature using far-UV circular dichroism. The CD spectra of the WT protein is typical for alpha-helical proteins with a minimum at 222nm (**Figure S1 A**). The thermal denaturation curves for all proteins measured at 222nm show a cooperative transition from the folded to the unfolded state (**Figure S1 B**). The melting temperatures (Tms) for the G408E, L540S and L808F mutant proteins are very similar to the WT protein (~ 39°C) and the Tms for the F711S, E767K mutants have a slightly lower Tms (~ 36°C) (**Figure 5A**).

**Figure 5.**
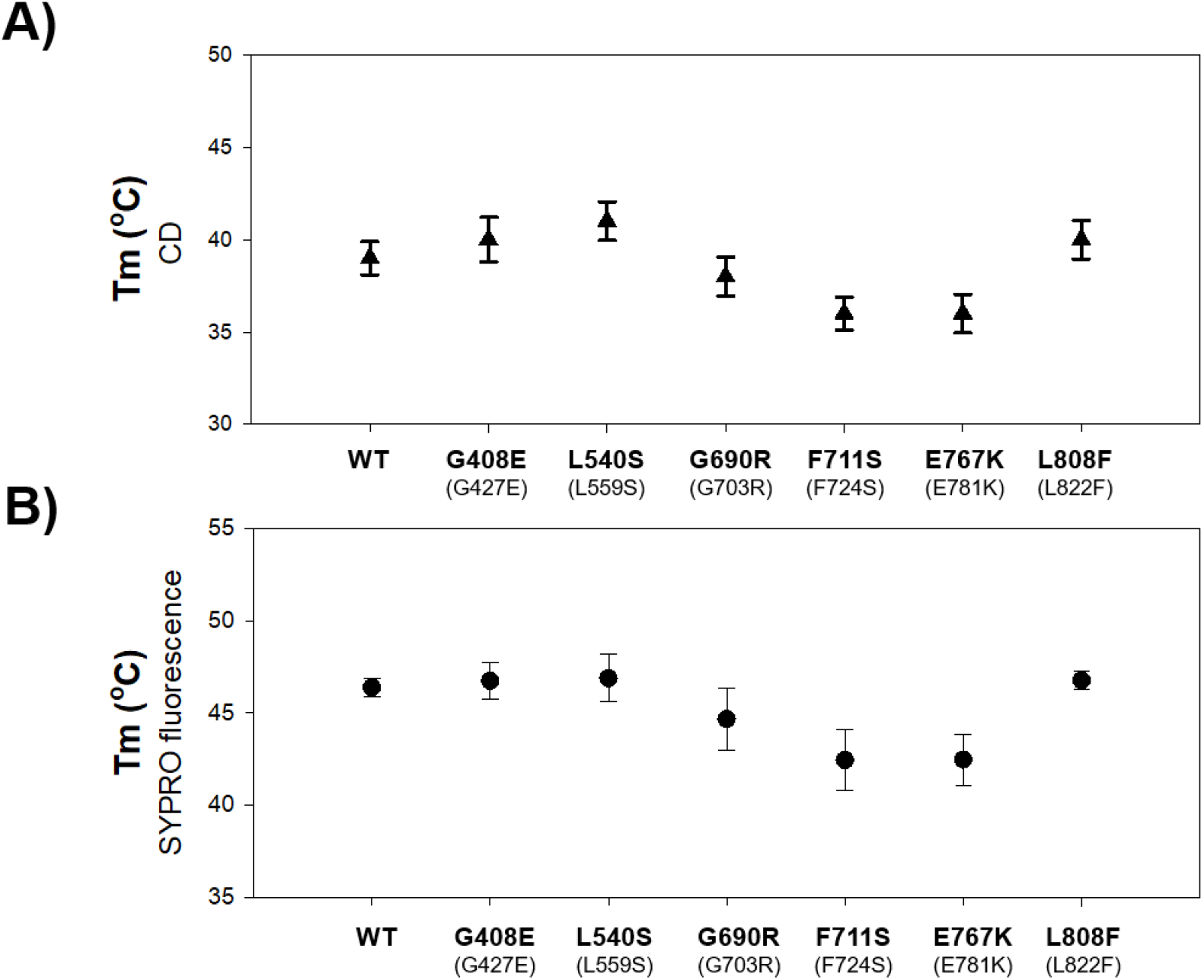
Temperature-sensitive mutations have a small effect on overall thermal stability of the UNC-45B chaperone. **A)** Melting temperatures of the UNC-45B proteins determined by CD spectrososcopy. Proteins at 1 uM were heated from 5 to 60oC, and their denaturation was followed by CD ellipticity at 222 nm. Each point in the measured Tm represent two measurements per temperature point (at 222 nm), and these were repeated two times (n=4). **B)** Melting temperatures of the UNC-45B proteins determined by DSF fluorescence spectrososcopy using the SYPRO Orange dye. Six DSF melting curves were obtained to determine the Tm for each protein (n=6 for each data point). Error bars are +/− SEM. The equivalent *ts* mutations in the *C. elegans* UNC-45 sequence are shown in parenthesis.

As an independent assay to quantify the effects of the mutations on UNC-45B thermal stability, we used differential scanning fluorimetry (DSF) with the protein stain SYPRO Orange to determine their melting temperatures. The stain intercalates with gradually exposed hydrophobic residues on the protein’s surface, allowing for melting temperature estimation based on the half-maximal SYPRO fluorescence in a Boltzmann approximation (23) (**Figure S2**). A major advantage of using fluorescence for protein thermal unfolding rather than traditional CD spectroscopic methods is that it can be done with greater speed, and thus with assaying many more samples and more robust statistics are obtained (23). Also this method is more sensitive to global exposure of hydrophobics, rather than local secondary structure changes that occur during thermal unfolding. As the data show (**Figure 5B**), the overall thermal stability of the G408E, L540S and L808 mutants, as determined by their Tm, is similar to the WT (~ 46°C) with the exception of the F711S and E767K mutants that show a slightly lower Tm (~ 43°C).

### Measuring chaperone activity of mutant UCS proteins using a S1 Mg-ATPase heat inactivation protection assay

To evaluate the effect of these mutations on the chaperone activity of the UNC-45B mutants, we measured how effective these proteins are in protecting the myosin heads (myosin subfragment 1, S1) from unfolding (or partial unfolding) when exposed to heat stress. We assessed unfolding under heat-shock conditions by determining the effect of elevated temperatures on its Mg-activated ATPase activity. As a positive control, we used the full length WT UNC-45B protein (20). **Figure 6** shows the relative Mg-ATPase activity of S1 + UNC-45B for all six mutants. As the data show, all mutants had a significantly lower heat protection effect on ATPase activity (range from ~70% for F711S to ~80% for L808F).

**Figure 6.**
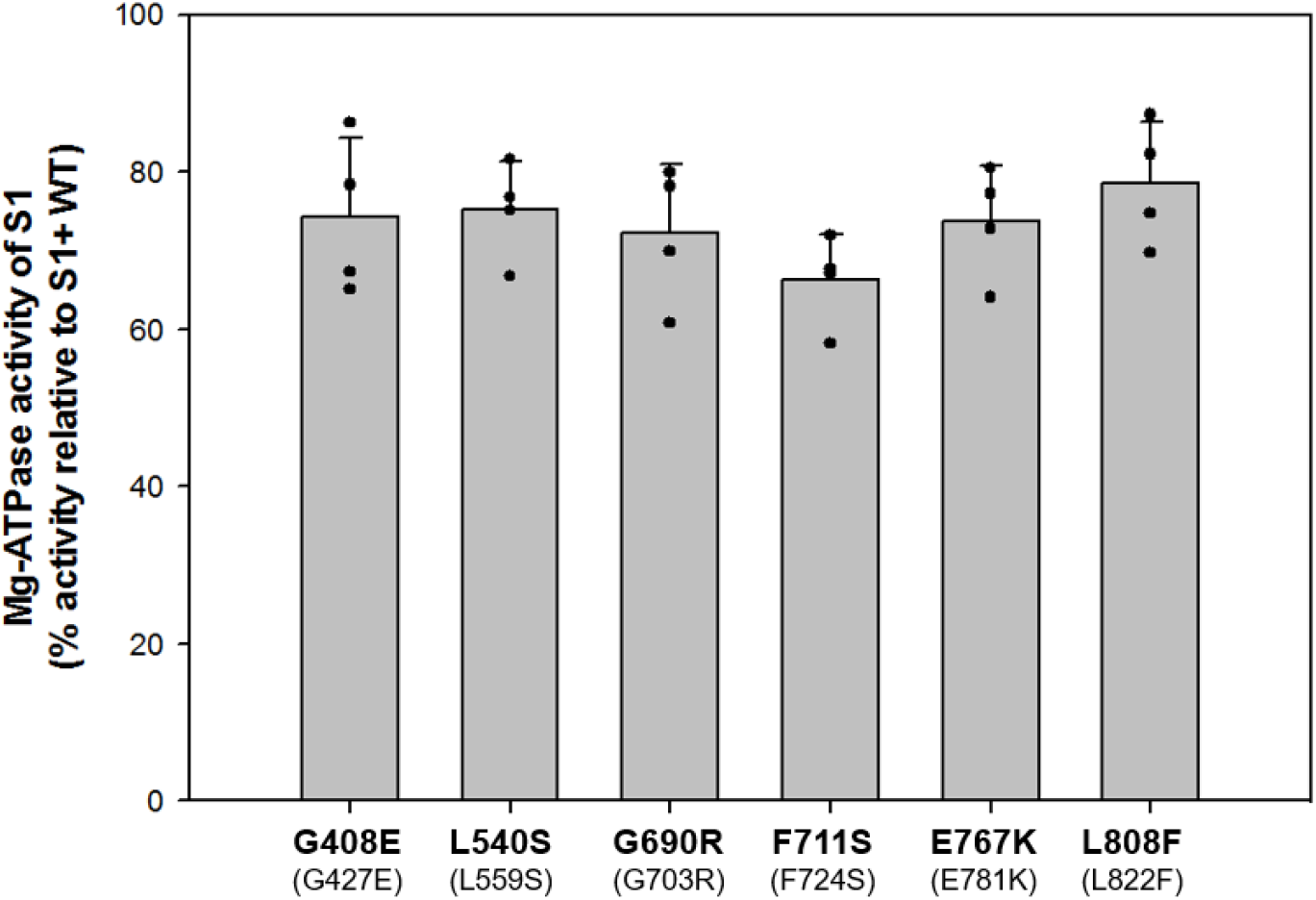
All 6 ts UNC-45B mutant proteins have significantly reduced ability to chaperone myosin as measured by a myosin S1 Mg-ATPase heat inactivation protection assay. The S1 Mg-ATPase activity is plotted as the % activity relative to UNC-45B wild-type (set to 100%). Assays were run in duplicates in two independent separate experiments (n=4 for each protein). The equivalent *ts* mutations in the *C. elegans* UNC-45 sequence are shown in parenthesis.

### Discussion and Conclusions

Temperature-sensitive (ts) mutants of a gene are ones in which there is a marked drop in the level or activity of the encoded protein when the gene is expressed at the restrictive temperature. At the lower permissive temperature, the phenotype of the mutant is very similar to that of the WT. Ts-mutants provide an extremely powerful tool for studying protein function in model organisms like *S. cerevisiae*, *C. elegans,* and *Drosophila* which do not maintain their internal temperatures and were instrumental in identifying the *unc-45* gene in *C. elegans* (5). However, the molecular mechanisms underlying temperature sensitivity have remained elusive. In the case of the myosin chaperone UNC-45, these mutations may alter protein stability, chaperone activity/interactions with myosin, or both. After first identifying two new *unc-45* ts mutants in the chaperone UCS domain, namely, G703R and F724S, we assessed the steady state levels of UNC-45 in a total of 6 *unc-45* ts missense mutants. Except for one of them (F724S), they show decreased levels of UNC-45 protein (ranging from 40% to 80% reduction) compared to wild type (**Figure 1D**). The total of six ts-mutations, G427E, L559S, G703R, F724S, E781K and L822F all are located in highly conserved regions (**Figure 3; Table 1**), and are located at residues that are conserved between nematode UNC-45 and human UNC-45B. *E. coli* was used to express and purify human UNC-45B with the comparable mutations. These amino acid changes in UNC-45B do not have a large impact on its thermal stability as measured by CD and DSF spectroscopy (**Figure 5**). Two ts-mutations (i.e. F711S and E767K) slightly decrease the thermal stability of UNC-45B by about 3°C (**Figure 5**). Interestingly, these two mutations are located in the center of the UCS domain (helices 13H1 and 14H2, respectively) and nearby to the ‘‘putative myosin-binding’’ groove region, helix 13H3 (20). We speculate that these two mutations may destabilize the compact structure of the groove region leading to a local structural change and a small overall destabilization. However, all the mutants exhibit a significantly lower chaperoning activity on protecting myosin heads from heat damage (**Figure 6**), including F711S, which in the nematode as F724S, does not affect its stability *in vivo*. Interestingly, in *C. elegans*, UNC-45 F724S is exceptional in that it shows increased levels of UNC-45 protein (**Figure 1D**) and its client myosin MHC B (**Figure 2B**) despite having increased myosin aggregation and disorganization of assembled thick filaments (**Figure 1A**) at the restrictive temperature. The comparable F711S mutant UNC-45B in vitro is one of the two worst UNC-45B proteins in terms of thermal stability (**Figure 5**), and shows the lowest myosin chaperone activity (**Figure 6**). These results on UNC-45 F724S and UNC-45B F711S, demonstrate that increased levels of a dysfunctional protein are just as deleterious as reduced levels of a dysfunctional protein. However, results on UNC-45 F724S are in contrast to transgenic overexpression of wild type UNC-45, which results in reduced assembled thick filaments and reduced levels of MHC B (19).

In terms of biochemical activities, our results are basically consistent with those by Hellerschmied et al., 2019 who found that four ts-mutations G427E, L559S, E781K and L822F have the same or even slightly higher thermal stability than the *C. elegans* WT UNC-45 chaperone (17). Using a cellular chaperone assay that measures the production of soluble myosin in insects cells, they found that these four UNC-45 ts-mutations had a negative effect on the myosin folding activity. Overall, our results indicate that the ts-mutations in UNC-45 lead to reduced and abnormal thick filament assembly by a combination of reduced UNC-45 protein stability and reduced UNC-45 chaperoning function. One potential reason for a difference in *in vivo* vs. *in vitro* mutant protein stability is that the mutations result in a change of post translational modifications that ultimately lead to increased degradation. This could occur by altering UNC-45’s interaction with another protein that normally covers a PTM site or causing residues to be slightly more or less exposed for modification. This may provide evidence that these conserved residues not only play a role in chaperoning activity, but also *in vivo* UNC-45 stability and/or regulation.

## MATERIALS AND METHODS

### *Caenorhabditis elegans* strains

Standard growth conditions for *C. elegans* were used (24). Wild-type nematodes were the N2 (Bristol) strain. The following strains were used in this study: *unc-45(e286), unc-45(m94), unc-45(b131), unc-45(su2002), unc-45(gk576605), unc-45(gk615477), and unc-45:mNeonGreen. e286*, *m94*, *b131* and *su2002* were obtained from the *Caenorhabditis* Genetics Center. VC40328 (*unc-45(gk576605)*, G703R) and VC40392 (*unc-45(gk615477)*, F724S) were 2 of 6 strains obtained from the million mutation project (MMP) (18) that had missense mutations in conserved residues of the UCS domain and showed disorganized thick filaments when grown at 25°C. They were outcrossed with N2 wild type five times to eliminate most of the background mutations of a typical MMP strain, to create strain GB337 for *unc-45(gk576605)* [G703R], and strain GB338 for *unc-45(gk615477)* [F724S]. The strain PHX789, *unc-45(syb789*), is a CRISPR generated strain which expresses UNC-45 with a C-terminal mNeonGreen tag. PHX789 was created by SunyBiotech (http://www.sunybiotech.com). PHX789 was outcrossed 2X to wild type to generate strain GB319.

### Generation of antibodies to UNC-45 and Western Blots

Glutathione S-transferase (GST) and MBP fusions of the 123 C-terminal residues of UNC-45 were expressed in *E. coli* after cloning into pGEX-KK1 and pMAL-KK1. The GST fusion protein was supplied to Noble Life Sciences (Woodbine, Maryland) for production of rabbit antibodies. Anti–UNC-45 was affinity-purified using Affigel (BioRad)-conjugated to the MBP fusion, as described previously (25). The method of Hannak et al. (26) was used to prepare toal protein lysates from wild type and the 6 *unc-45* mutant strains. Equal amounts of total protein from these strains were separated by 12% or 4-15% polyacrylamide-SDS Laemmli gels, transferred to nitrocellulose membranes, and the top half was reacted with affinity-purified anti–UNC-45 at 1:5,000 or anti-MHC B (5-13; (8); gift of Henry F. Epstein, now deceased) at 1:50,000, and the bottom half was reacted against a rabbit polyclonal antibody directed to human histone H3 (Abcam, cat. no. ab1791) at 1:40,000, and then with goat anti-rabbit immunoglobulin G conjugated to HRP at 1:10,000 and visualized by ECL. The quantitation of steady-state levels of protein was performed as described in Miller et al. (27). The relative amount of each of these proteins in each lane was normalized to the amount of Histone H3.

### Immunofluorescent staining of body wall muscle

Adult nematodes were fixed and immunostained using the method described by Nonet et al. (28) with further details provided by Wilson et al. (29). The primary antibody, anti–myosin heavy chain A (MHC A; mouse monoclonal 5-6; obtained from the University of Iowa Hybridoma Bank) (8), was use at 1:100 and the secondary antibody, anti-mouse Alexa 594 (Invitrogen), at 1:200. Images were captured at room temperature with a Zeiss confocal system (LSM510) equipped with an Axiovert 100M microscope and an Apochromat ×63/1.4 numerical aperture oil immersion objective. The color balances of the images were adjusted by using Photoshop (Adobe, San Jose, CA).

### Human UNC-45B Protein Structure Modelling and Chimera Structural Analysis

The homology model of the human UNC-45 protein structure was done using Phyre2 (30) and SWISS-MODEL (31) using the *C. elegans* 3D structure (PDB ID: 4i2z) as a reference crystal structure. The modeled human UNC-45B protein structure was displayed with Chimera (22) and single amino acid mutations were inserted using the rotamer tool and then energy minimized to minimize interatomic clashes and contacts based on van der Waals radii.

### Mutagenesis, Protein Expression and Purification of proteins

UNC-45B (wild-type and mutants) from *Homo sapiens* was codon optimized for expression in *Escherichia coli*, synthesized (GenScript, Piscataway, NJ) and subcloned into a pET-28A vector (EMD Millipore, Billerica, MA). Mutations were introduced into UNC-45B wild-type using the Q5® Site-Directed Mutagenesis Kit (NEB). Recombinant protein expression was induced in BL21 DE3 when optical density (OD600) reached 1-1.2 with 0.02 mM IPTG for 16 h at 15 °C. Harvested cells were resuspended in lysis buffer (50 mM Tris-HCl, 50 mM NaCl, 40 mM imidazole, 2 mM TCEP, 10% glycerol, pH 8.0) containing 1 mg/ml lysozyme and incubated for 30 min at RT. To promote lysis cells further samples were sonicated 6 × 20 s with 20 s breaks (200 – 300 W) on ice. To break down DNA and reduce viscosity, 1 μl of Universal nuclease (Pierce; 125U/L of culture) and 10 μl of DNaseI (NEB; 20U/L of culture) were added and the mixture allowed to incubate for 10 min at RT. Insoluble material was removed by centrifugation at 30,000 x g for 30 min at 4 °C and supernatant was filtered through 0.45 μm syringe filter afterwards. Supernatant was incubated with lysis buffer equilibrated HisPur Ni-NTA Resin (Thermo Scientific) and incubated overnight at 4°C. Resin with bound proteins were collected by centrifugation at low force (800 x g for 2-3 min), washed by 4 washes with ice cold wash buffer (50 mM Tris-HCl, 50 mM NaCl, 60 mM imidazole, 2 mM TCEP, 10% glycerol, pH 8.0) and with last wash applied to the column cartridge. His-tagged recombinant proteins were eluted in 1 ml fractions with elution buffer (50 mM Tris-HCl, 50 mM NaCl, 250 mM imidazole, 2 mM TCEP, 10% glycerol, pH 8.0) and immediately 2 mM of EDTA pH 8.0 were added. Eluted fractions containing significant portion of recombinant protein were pooled and dialyzed against storage buffer (50 mM Tris-HCl, 50 mM NaCl, 2 mM EDTA, 2 mM TCEP, 10% glycerol, pH 8.0) at 4°C. Proteins were concentrated to 20-40 μM, flash frozen and stored at −80 °C.

### Protein Thermal Stabilty using CD Spectroscopy

Far UV CD experiments were done as previously described (20,32). Briefly, spectra (1990-250nm) of UNC-45B proteins at 1 μM (in 30 mM phosphate buffer pH 7.4, 100 mM KCl, 1 mM MgCl_2_, 1 mM TCEP) were recorded on a Jasco J-815 Spectrometer. For determination of the melting curves, signals in the far-UV CD region (222 nm) were monitored as a function of temperature.

### Protein Thermal Stabilty using Fluorescence Spectroscopy

Differential scanning fluorimetry (DSF), also known as ThermoFluor or Thermal Shift was done basically as described in (23). All proteins were used at a final concentration of 1μM for this assay. SYPRO Orange (Invitrogen S6651) was used at a final concentration of 5x (based on the 5000x stock solution as supplied by Invitrogen) in potassium phosphate buffer (in 30 mM phosphate buffer pH 7.4, 100 mM KCl, 1 mM MgCl_2_, 1 mM TCEP). All DSF experiments were carried out with StepOnePlus™ Real-Time PCR System (Thermo Fisher Scientific). Each sample was divided to three 30 μL replicates. Sample solutions were dispensed into a MicroAmp® Fact Optical 96-well reaction plate (Thermo Fisher Scientific) and the plate was sealed with MicroAmp® Optical Adhesive Film (Thermo Fisher Scientific). Fluorescence intensity was measured using 580 nm excitation and 623 nm emission filters. Controls included samples containing buffer only and dye plus buffer only. Temperature was continuously increased from 20°C to 99°C at 0.5 °C/min. Melting curves were directly exported from the instrument, and then were analyzed with the Thermo Fisher software (TM) Software v1.0. Six melting curves were obtained for each protein.

### S1 Mg-ATPase heat inactivation protection assay

Protection of myosin enzymatic activity from heat by UNC-45B proteins was essentially done as previously described (20). Briefly, 0.4 μM of S1 Fragment (Rabbit Skeletal Muscle, Cytoskeleton #CS-MYS04) was incubated either alone or with 2 μM UNC-45B proteins for 10 minutes either at 42°C or on ice in Mg-buffer (10 mM PIPES-KOH, 100 mM KCl, 10mM MgCl_2_, 0.1mM CaCl_2_, 0.3 mM EGTA, 2 mM TCEP, pH 7.4). Heat exposed samples were cooled down on ice for 1 minute and mixed 1:1 with 1 mM ATP in Mg-buffer to initiate ATP hydrolysis for 15 min at room temperature. S1 ATPase activity was quenched with PiColorLock™ Gold Reagent (Expedeon Inc., San Diego CA 92121) and free Pi detection was carried out according to the manufacturer’s recommendation. Samples were read at wavelength 620 nm using a FLUOstar Optima Microplate reader (BMG Labtech Inc.). Readings were blank corrected and used to calculate relative activity to S1. The assay was carried out in duplicate in at least two independent experiments (n = 4 for each protein).

#### Statistical analyses

Unless otherwise stated, data are reported as mean ± standard error of the mean. Western Blot statistical analyses were made using GraphPad Prism Software (version 4.0). Comparisons of three or more means used one-way ANOVA and Bonferroni-adjusted unpaired *t* tests. Statistical significance was assigned as not-significant for P > 0.05, * for P ≤ 0.05 and *** for P ≤ 0.001.

## ACKNOWLEDGMENTS

This work was supported by National Institutes of Health grant R01GM118534 to G.M.B. and A.F.O. We thank Drs. Luis Holthauzen and Pawel Bujalowski for their invaluable help with the CD experiments.

## Author Contributions

G.M.B. and A.F.O. conceived the study; I.G. and T.M. expressed and purified proteins; T.M. performed CD and analyzed data with A.F.O.; T.M. performed chaperone experiments and analyzed data with A.F.O.; J.M. performed the SYPRO Orange experiments and analyzed data with A.F.O.; C.J.M. and H.Q. performed worm immunotaining, generation and evaluation of antibodies, and western blots and analyzed the data with G.M.B.; G.M.B., A.F.O. and C.J.M., wrote the manuscript.

---

**Figure S1:**
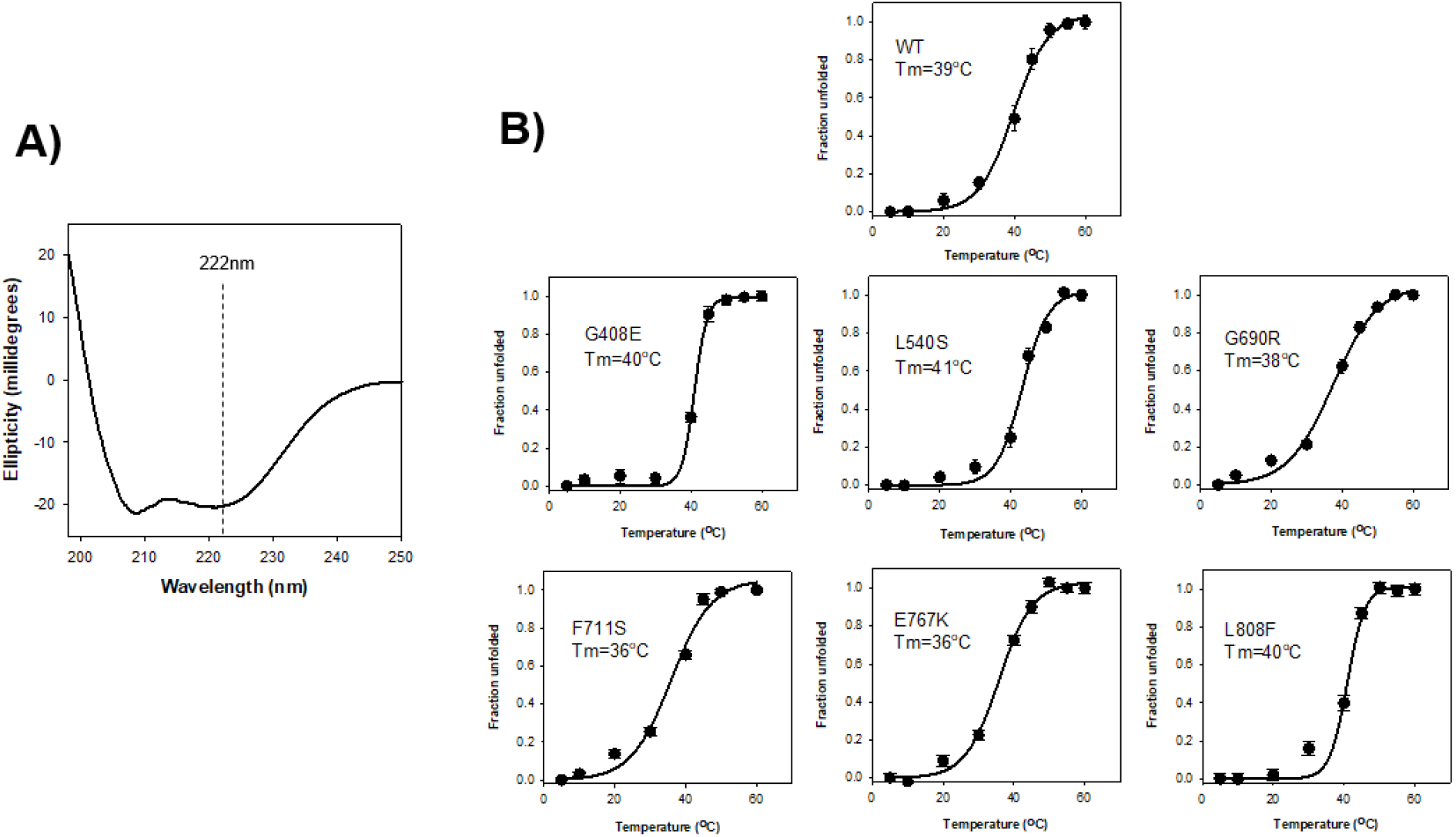
Circular dichroism analysis of the WT and mutant UNC-45B proteins. **A)** A typical Far‐UV circular dichroism of the WT UNC-45B protein. The protein concentration was 1 μM in 30 mM phosphate buffer pH 7.4, 100 mM KCl, 1 mM MgCl2, 1 mM TCEP buffer. A 0.1 cm path length cuvette was used. The % of a-helical content was estimated 58% for the WT using the Bestsel tool (http://bestsel.elte.hu.). **B)** Thermal denaturation of WT and mutant UNC-45B proteins monitored by circular dichroism. UCS proteins at 1uM were heated from 20 to 65 °C, and their denaturation was followed by circular dichroism ellipticity at 222 nm.

**Figure S2.**
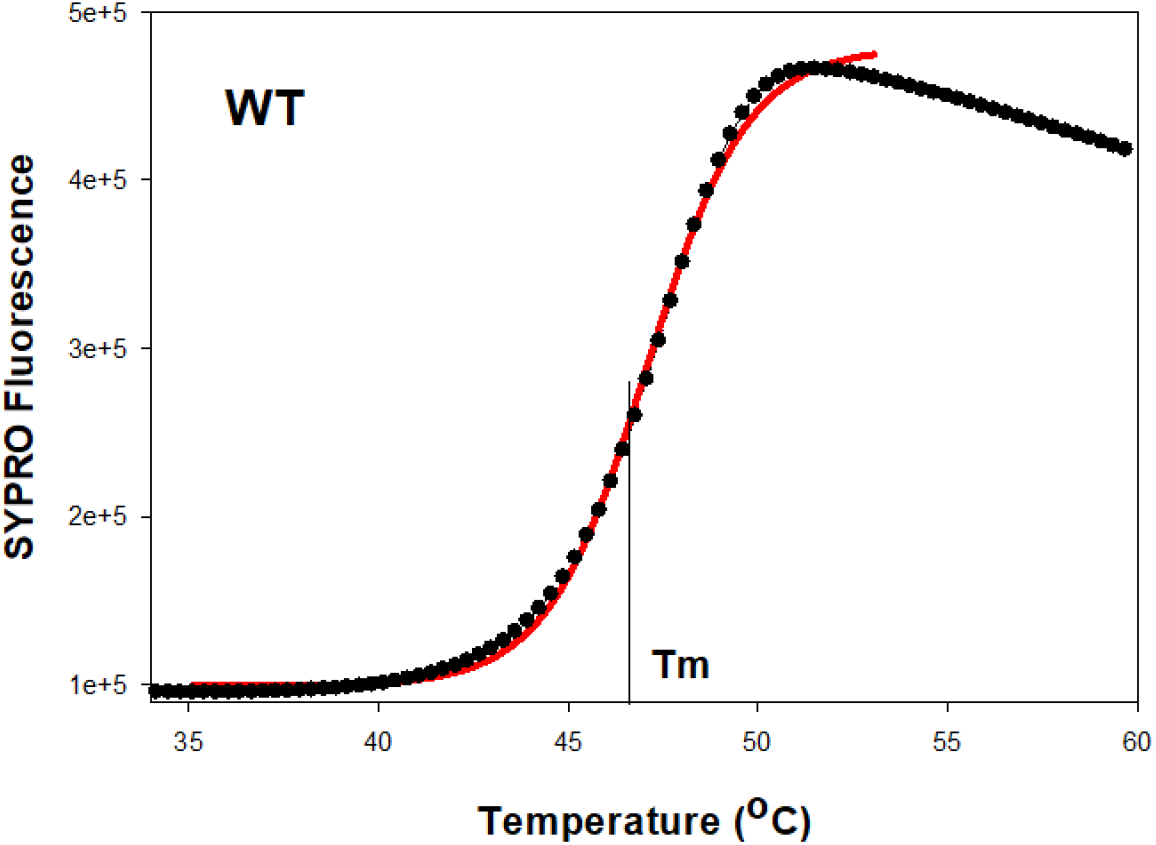
Typical thermal denaturation profile of a WT UNC-45B sample using Differential scanning fluorimetry (DSF). Fluorescence emission changes with the temperature. The sigmoidal curve indicates the cooperative unfolding status of the UNC-45B protein from trace amounts of SYPRO Orange bound to the native protein. The peak indicates that all proteins are unfolded to linear peptides or that the hydrophobic core is exposed to SYPRO Orange (23). The midpoint of the transition curve is the melting temperature (*T*m). The red line corresponds to a fit to a sigmoidal curve.

